# The Effects of Ribosome Autocatalysis and Negative Feedback in Resource Competition

**DOI:** 10.1101/042127

**Authors:** Fiona A Chandra, Domitilla Del Vecchio

**Affiliations:** Department of Mechanical Engineering, Massachusetts Institute of Technology 77 Massachusetts Ave 3-469A Cambridge, MA 02139

**Keywords:** Resource competition, ribosome, autocatalysis

## Abstract

**Background:** Resource competition, and primarily competition for ribosomes, can lead to unexpected behavior of genetic circuits and has recently gained renewed attention with both experimental and theoretical studies. Current models studying the effects of resource competition assume a constant production of ribosomes and these models describe the experimental results well. However, ribosomes are also autocatalytic since they are partially made of protein and autocatalysis has been shown to have detrimental effects on a system’s stability and robustness. Additionally, there are known feedback regulations on ribosome synthesis such as inhibition of rRNA synthesis via ppGpp.

**Results:** Here, we develop two-state models of ribosome and protein synthesis incorporating autocatalysis and feedback to investigate conditions under which these regulatory actions have a significant effect in situations of increased ribosome demand. Our modeling results indicate that for sufficiently low demand, defined by the mRNA level of synthetic genes, autocatalysis has little or no effect. However, beyond a certain demand level, the system goes through a transcritical bifurcation at which the only non-negative steady state is at zero ribosome concentration. The presence of negative feedback, in turn, can shift this point to higher demand values, thus restoring the qualitative behavior observed in a model with a constant ribosome production at low demand. However, autocatalysis affects the dynamics of the system and can lead to an overshoot in the temporal response of the synthetic genes to changes in induction level.

**Conclusion:** Our results show that ribosome autocatalysis has a significant effect on the system robustness to increases in ribosome demand, however the existing negative feedback on ribosome production compensates for the effects of the necessary autocatalytic loop and restores the behavior seen in the system with constant ribosome production. These findings explain why previous models with constant ribosome production reproduce the steady state behavior well, however incorporating autocatalysis and feedback is needed to capture the transient behavior.

## Background

Ribosomes as a limited resource have gained renewed attention lately with numerous studies on resource limitations in synthetic biology (1, 2, 3, 4). Ribosomes play a key role in gene expression for both synthetic plasmid and host genes. In (5, 6, 7), the authors review the importance of host cell-dependent factors in synthetic biology design, including the effects of limited cellular resources in the expression of synthetic and host genes. The expression of a constitutive gene or an endogenous gene may unexpectedly decrease when the expression of another gene is increased due to sequestration of common resources such as RNA polymerases and ribosomes, as shown experimentally between two plasmid genes in (1). In (3), Ceroni et al. uses a chromosomal insertion of green fluorescent protein (GFP) to monitor gene expression capacity to show that ribosome demand from synthetic constructs also affects chromosomal gene expression, at least in the short term. While RNA polymerase is also a limited resource (8), it has been shown that ribosomes present the main bottleneck both in protein synthesis and in cell growth (1, 9, 4).

Resource competition has also been studied using computational models. Qian et al. showed mathematically that resource competition can lead to hidden interactions that explain previous experimental results (2). When two seemingly unrelated genes compete for the same resources, the expression of one can lead to a decrease of expression in the other, leading to an effect equivalent to repression. Similar effects are also seen in cell-free systems (10). In (17), Algar et al used Ribosome Flow Model of translational elongation to show the importance of ribosome sequestration in gene expression and provide insight on synthetic construct design to mitigate such effects (17). In (16) the authors present a framework to calculate a network’s sensitivity to competition using response coefficients from metabolic control analysis (18). Most models of resource competition assume that ribosomes are present at a constant total level in the cell (1, 2, 10 and 17). These models predict the effects of resource competition behavior quite well; however, we know that ribosomes are produced in an autocatalytic manner, which does not guarantee a constant total ribosome concentration. In *E. coli*, ribosomes are composed of 55 protein subunits which make up 30% of ribosomes by weight (20). Ribosomes must therefore be translated by ribosomes themselves, resulting in an autocatalytic loop, and total ribosome concentration may change when the amount of mRNA to be translated changes. In addition, the autocatalysis adds to the resource competition problem since ribosomes also compete with cellular or synthetic plasmid genes for the same pool of resources. This could lead to a drop in the total ribosome concentration as synthetic gene expression is increased.

Autocatalysis has been shown to have detrimental effects in the robustness of a system to perturbations both in the steady state and temporal responses (25). Computational models of protein translation incorporating ribosome autocatalysis have been analyzed to determine the optimal ribosome concentration and allocation for maximal growth (18), while (14) looks at a phenomenological model to study how the cell allocates resources in different growth conditions. Other studies include a more detailed model of ribosome flow, incorporating stepwise translation of individual codons (17). These models provide some insights on the effects of autocatalysis on cellular growth, but not on the effect of autocatalysis on ribosome competition in particular. In this paper, we compare three models of ribosome production to explore the effects of the ribosome autocatalysis on the steady state behavior of competing genes as well as on the system’s stability and its transient response to perturbations.

It is known that ribosome production is tightly regulated in the cell based on nutrient level and translational activity (23). When ribosome is bound to mRNA and encounters an uncharged tRNA, it activates an enzyme called RelA (bound to ribosomes), which then produces an intracellular signaling molecule, guanosine pentaphosphate (ppGpp). ppGpp then binds to a protein called DksA and this complex inhibits rRNA transcription (21, 24). When amino acid level is low, the levels of uncharged tRNA increases, however when translation activity is low, the uncharged tRNA is less likely to encounter RelA bound to ribosomes. When translation demand and ribosome levels are high, there is higher level of both RelA bound to ribosomes and of DksA, resulting in inhibition of rRNA. Thus, there is a negative feedback on the ribosome production. This feedback senses a combination of translational demand, ribosome level, and amino acid level in the cell. Other models such as (12, 15) incorporates the feedforward regulation from amino acid pool to ribosome synthesis while in this paper we focus on the feedback loop from ribosome concentration. In (11, 33), the authors presented a model of protein translation and cell growth with ppGpp feedback but this model does not incorporate ribosome autocatalysis, modeling ribosomes only as composed of ribosomal rRNA (rRNA). In (31), the authors discuss the effects of ppGpp feedback on growth rate and its consequence on gene expression. The growth rate of cells also initially decreases upon expressing a synthetic gene, but recovers after a few generations (11). The recovery is hypothesized to be due to the negative feedback discussed above. Here, we explore the interplay between autocatalysis and the ppGpp-dependent negative feedback on the expression of ribosome in the presence of synthetic mRNA.

Theoretical analysis and simulations of our mathematical models show that autocatalysis leads to a steeper decrease in free ribosome concentration in the presence of increasing synthetic demand of ribosomes. However, negative feedback restores the steepness of the drop in ribosome concentration to a level similar to the model incorporating constant ribosome production. The negative feedback thus compensates for the deleterious effects of ribosome autocatalysis. This suggests that previously reported resource competition models using a constant ribosome pool would predict similar change in free ribosome level as the model with autocatalysis and feedback within a range of synthetic mRNA level. The autocatalytic system undergoes a transcritical bifurcation where there is no positive steady state. A strong enough negative feedback can shift the bifurcation point beyond physiologically relevant range of synthetic mRNA level (demand) and thus maintain a stable ribosome steady state for high levels of synthetic mRNA. Autocatalysis also has a significant effect on the dynamics of the system and results in overshoots in the time response of the synthetic gene, but not in the free ribosome, as seen in experiments (1).

## Results

To understand the effects of autocatalysis and negative feedback, in this paper we compare three different one-state models of ribosome production, shown in Figure 1. The first model assumes a constant production of ribosome. The second model incorporates autocatalytic production of ribosomes, and the third adds a negative feedback regulation on the ribosome production. The models described in the following sections were obtained by reducing a mechanistic model including mass action binding kinetics between ribosomes and mRNA and protein translation by using quasi-steady state approximation on bound ribosome-mRNA complexes (see Supplementary Information Section A for details).

**Figure 1.**
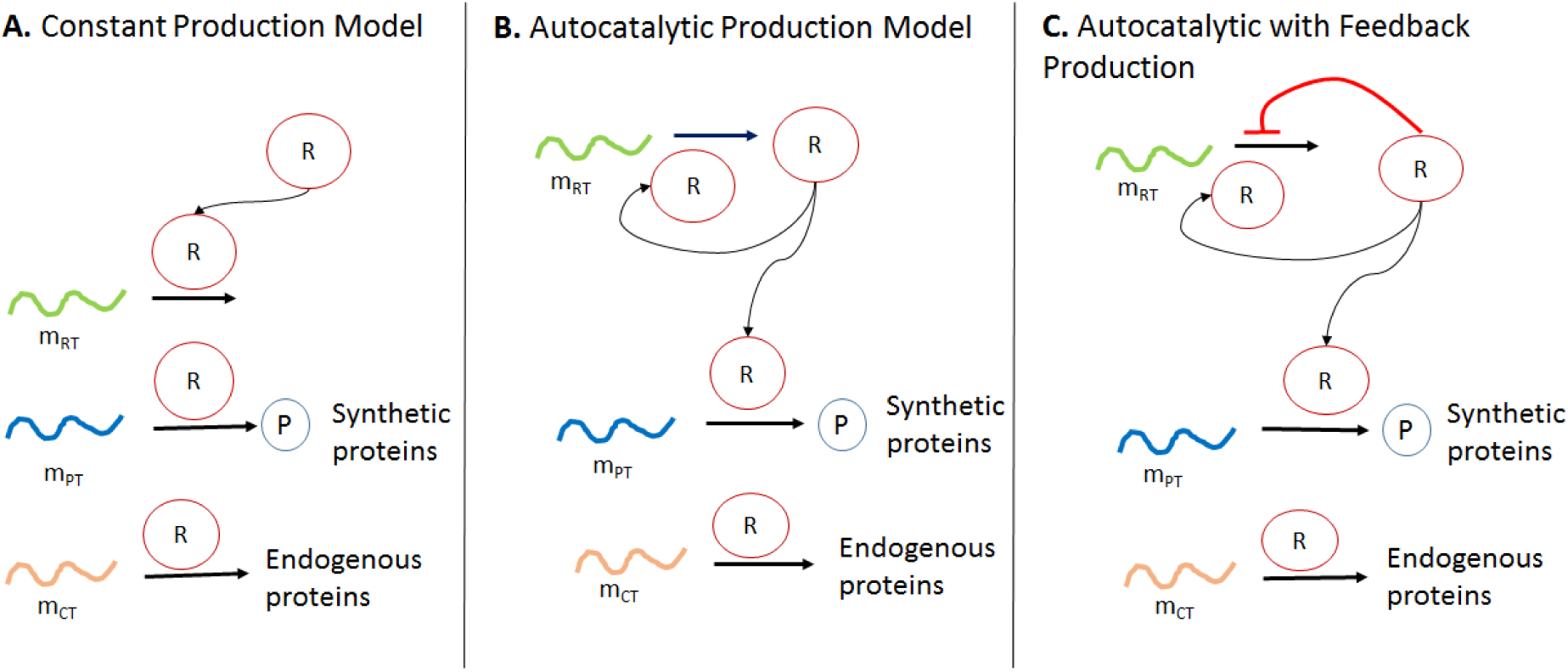
Three models of ribosome production. A) Model with constant ribosome *(R)* production. B) Autocatalytic model where ribosomes translate more ribosomes. C) Autocatalytic model with negative feedback where free ribosomes also inhibit the production of more ribosomes. Here, *mPT* is the mRNA of the inducible synthetic gene, *mRT* is the ribosomal mRNA, and *mCT* is the pool of all other mRNAs in the cell.

### A. Model with constant ribosome production

This model follows previous models with constant supply of ribosomes. Ribosomes (*R*) are produced at a constant rate *Vmax*, ensuring a constant steady state level of total ribosomes. Ribosomes are sequestered by the translation process of various mRNAs, including total ribosomal mRNA (*mRT*), which in the other two models will produce ribosomes, total mRNA of synthetic genes (*mPT*), and other mRNAs in the cell (*mCT*), which we assume to be constant for simplicity. Increasing *mPT* can be obtained experimentally by increasing either induction level of the synthetic gene or its copy number. To simplify the analytical derivations, we assume that the effective dissociation constant of ribosome binding to all mRNA types are the same. The main conclusions do not depend on this simplifying assumption as will be illustrated through simulation for the case where these values are different. The reduced order model (see SI Section A for model reduction) for ribosome concentration can thus be written as:

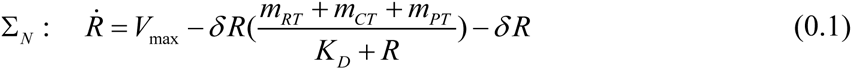

where δ is the dilution rate and *K*_*D*_ is the effective dissociation constant, which encompasses the binding of ribosomes to mRNA and dissociation both from spontaneous unbinding and translation completion.

The total ribosome concentration is the sum of the free ribosomes and the ribosomes bound to the different mRNAs in the cell (SI Section A):

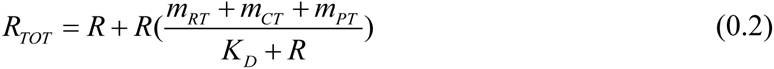

In Σ_N_, this is given by 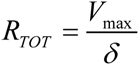 (see SI Section B).

The steady state concentration of free ribosomes can be obtained by setting *Ṙ* = 0 to determine how *R* depends on the demand *mPT*.

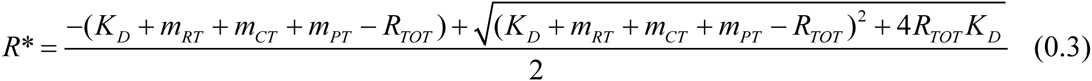

Free ribosome level *R* drops as *m*_*PT*_ is increased. The change in free ribosome concentration at a given *mPT* concentration is described by the following nonlinear equation:

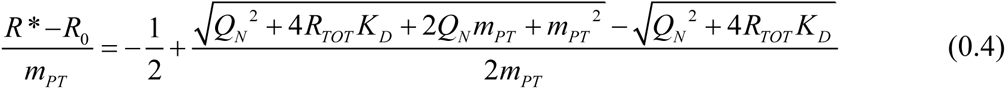

where *Q*_*N*_=*K*_*D*_+ *m*_*RT*_ + *m*_*CT*_ − *R*_*TOT*_ and *R*_0_ is the free ribosome concentration at *m*_*PT*_=*0* (see SI Section E).

For low enough *R*_*TOT*_, *Q*_*N*_ is positive and thus the second term on the right hand side is positive. When *RTOT* is too high, *Q*_*N*_ can be negative yet the second term can still be positive if *m*_*PT*_ is high. This implies that the free ribosome level drops faster at lower demand (when *mPT* is low) than at higher demand level. However, even when *Q*_*N*_ is negative, (1.4) is still bounded below by −1 (see Fig S4), therefore the steepest drop is bounded below by a line with slope −1.

For our simulation, we set *V*_*max*_ such that the total ribosome 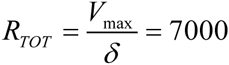, which is within the range of experimental observations (26). Total mRNA concentration in the cell *(mCT)* ranges from 3000-8000 copies per cell, depending on the growth rate (28). For our analysis, we have taken the average, which is *mCT*=5500 copies/cell. The ribosomal promoter activity ranges between 5-35% of the total promoter activity in the cell, depending on growth conditions (27). This suggests that the total ribosomal mRNA should be 5-35% of the total mRNA in the cell. In our simulations and analysis, *mRT* is chosen to be 10% of *mCT*, which lies within the observed range. Other parameter values and their references are described in Table 1. For these values, *Q*_*N*_ > 0, therefore while *R* drops nonlinearly with *m*_*PT*_, the steepest drop for this constant production model for high demand is bounded below by a line with slope −0.5. As *K*_*D*_ increases (indicating weaker ribosome binding site (RBS) strength), the drop in ribosome becomes less steep. This is consistent with the finding in (17) that stronger RBS (smaller *KD*) significantly decreases the robustness of ribosome level to increasing demand.

**Table 1.** Parameter values

| Parameter | Value<br>(references) | Description |
| --- | --- | --- |
| $k_t$ | 17.61 hr<sup>-1</sup> | Translation rate of ribosomal and cellular genes. Set such that $R_{TOT}=7000$ nm at $m_{PT}=0$ |
| $k_{tP}$ | 300 hr<sup>-1</sup> (2) | Translation rate of synthetic gene (i.e. GFP) |
| $\delta$ | 1 hr<sup>-1</sup> (2) | Dilution rate |
| $m_{RT}$ | 550 nm (27) | Total mRNA of ribosomes (10% of cellular mRNA) |
| $m_{PT}$ | 5-3000 nm | Total mRNA of synthetic gene (demand) |
| $m_{CT}$ | 5500 nm (28) | Total non-ribosomal cellular mRNA |
| $K_D$ | 1000 nm (1) | Effective dissociation constant of ribosome and mRNA complexes due to both unbinding and translation completion |
| $K_I$ | 1000-3000 nm | Effective inhibition constant |
| $R_{TOT}$ | 7000 nm (26) | Total Ribosome |
| $V_{max}$ | 7000 nm/hr | $V_{max} = R_{TOT}\delta$ |
| $V$ | Varies (nm) | Normalizing constant for ribosome production, varied to match total ribosome in cell $R_{TOT}=7000$ at $m_{PT}=0$. |

One of the main questions we would like to answer is how much demand can the cell withstand. In this model with constant ribosome production, there is always a stable steady state with *R*>*0*. 1. Fig 2A shows the change in free ribosome level as we increase the demand, modeled as the mRNA level of the synthetic gene *(mPT)*. The black line in Fig 2A shows the result for the constant production model using the parameter values described above. As previous studies suggest that ribosomes instead of RNA polymerase are the bottleneck for protein expression, for simplicity we assume that the other mRNA levels such as *mCT* and *mRT* remain constant (1, 4).

**Figure 2.**
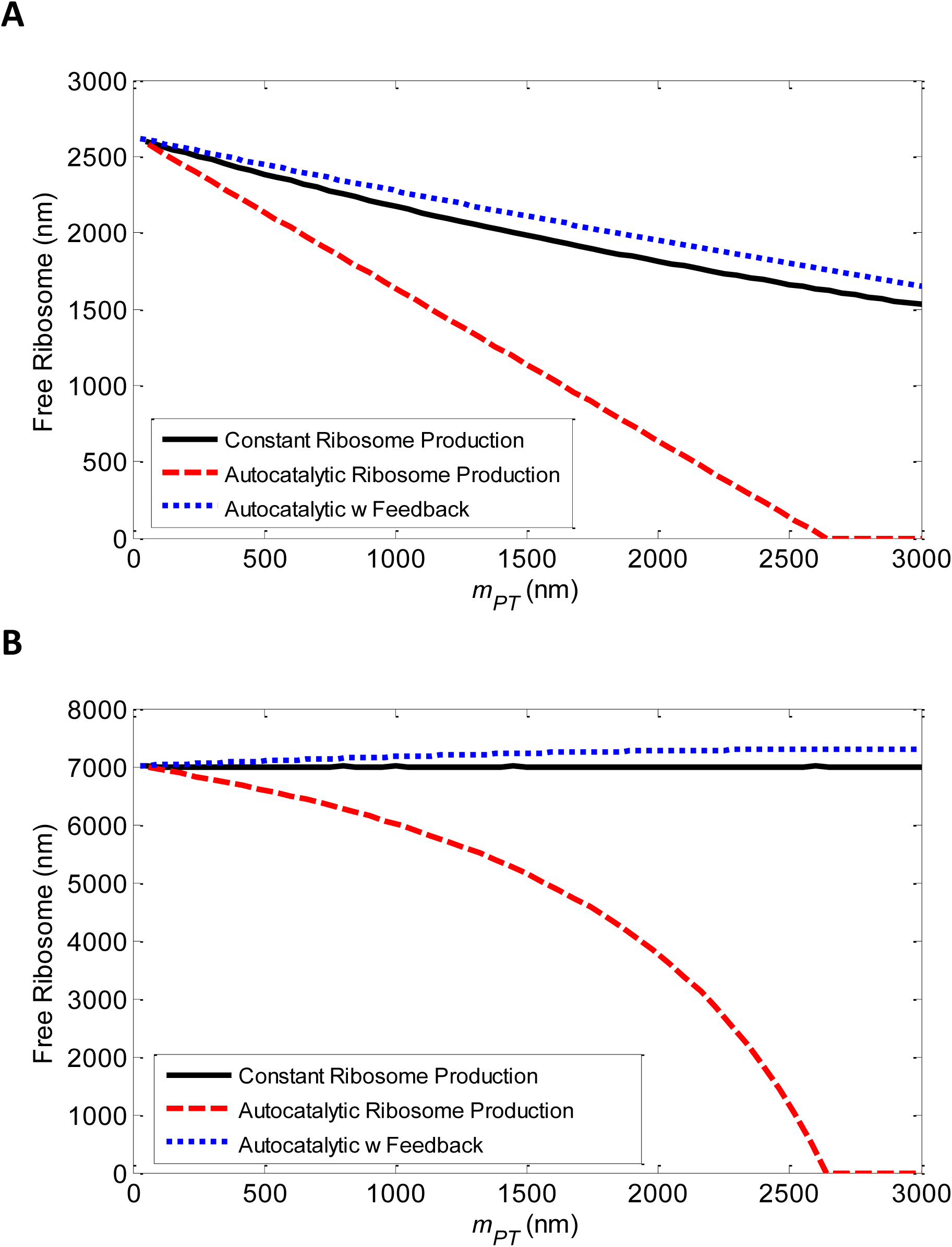
The change in free and total ribosome concentration as the demand *(mPT)* is increased for the three different models in Fig 1. A) The change in free ribosome level *(R)* as the demand *(mPT)* is increased. The autocatalytic system without feedback shows a steeper drop in the free ribosome level and eventually goes through a bifurcation and the steady state becomes *R*=*0*. B) The change in total ribosome level (*R*_*TOT*_) as *mPT is* increased. The total ribosome level at *mPT=0* for all models are set to *R*_*TOT*_=7000 nm, *m*_*RT*_=550 nm, *m*_*CT*_=5500 nm, *k*_*t*_=17.61 hr-^1^, *K*_*D*_=1000 nm, δ=1 hr^−1^. For the constant production system *Vmax*=7000. For the autocatalytic with feedback model, *K*_*I*_=3000 nm, *V*=5621.8 nm. Note that *Q*_*N*_ > 0 with these values.

### B. Model with ribosome autocatalysis

In this model, ribosomes translate ribosomal mRNAs *(mRT)* into more ribosomes. We assume that the ribosome translation rate follows a Michaelis-Menten form:

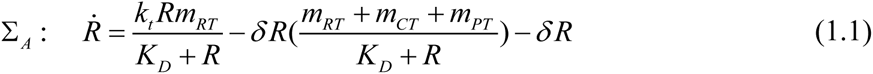

where *k*_*t*_ is the translation rate constant.

There are two steady state solutions to the above equation, given by:

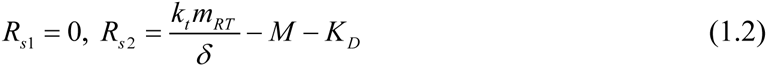

where *M* = *m*_*RT*_ + *m*_*CT*_ + *m*_*PT*_. For small *m*_*PT*_ (where 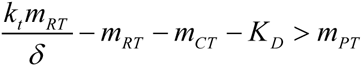),*R*_*S2*_>0 and is stable while *Rs1*=*0* is an unstable steady state (see SI Section D for details). When *mPT* increases to a high enough level, the system undergoes a transcritical bifurcation where the *R*=*0* steady state becomes stable and the other steady state at *R*<*0* becomes unstable (see SI Section D). In practice, depletion of ribosomes in the cell will result in cell death.

We determine from (2.2) the change in free ribosome concentration and find that in this case it decreases linearly with *mPT*, with slope given by (see SI Section E):

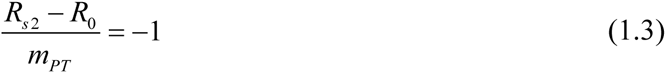

While the slope in this case doesn’t change with parameters, the autocatalytic system undergoes a bifurcation and the bifurcation point is parameter-dependent. In particular, beyond a certain demand, there is no positive non-zero steady state, there is only a steady state at *R*=*0*. We can analytically solve for the bifurcation point and find that there is a positive steady state *R*>*0* (which is stable, see SI Section D for details) when:

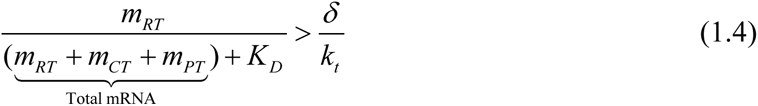

While Eq (2.3) shows that the slope does not change, Eq (2.4) shows that the bifurcation point is parameter dependent and moves to higher demand *mPT* as the dissociation constant *KD* is increased (see Fig 3). To estimate the demand or *mPT* from a high copy number plasmid, we assume that 1 nm = 1 molecule of mRNA and note that a high copy number plasmid such as pUC can have 500-700 copy number (29). Assuming an average of 2-3 mRNA transcripts per gene copy for highly expressed genes (30), the external demand given by a high copy plasmid would be around 1000-2100 mRNA copies. Using parameter values taken from literature (see Table 1), we see in Fig 3 that the bifurcation point for a range of *K*_*D*_ is within the range of gene expression from a high copy plasmid, indicating that without feedback the autocatalytic system is not robust enough to prevent the free ribosome concentration to collapse to zero. In Fig 2, we see that the autocatalysis introduces a significant drop in the free ribosome level as the demand is increased. The drop is much steeper in the autocatalytic case than in the constant ribosome production case, indicating that autocatalysis reduces the robustness of free ribosome concentration to external demand.

**Figure 3.**
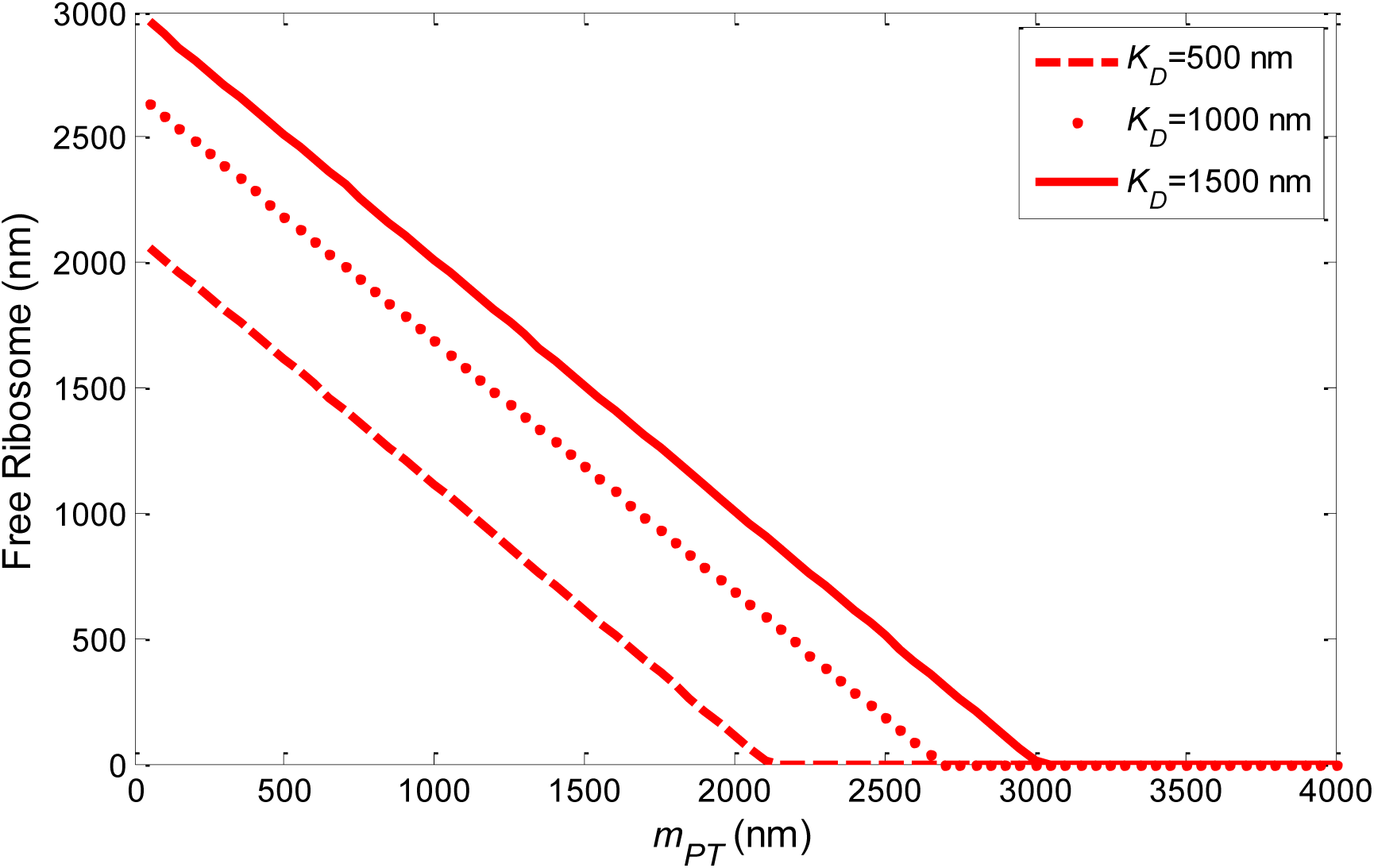
The change in ribosome concentration as *mPT* is increased for different values of *KD* in the autocatalytic system with no feedback Σ_A_. At lower *K*_*D*_ (stronger RBS), the bifurcation point occurs at lower demand *mPT*. The slope for all *K*_*D*_ values remains the same.

The bifurcation point solution in (2.4) shows that the minimum ribosomal to total mRNA ratio required for a stable *R*>*0* steady state increases with the dilution rate δ. This suggests that as growth rate increases (which would increase the dilution rate), the cell must produce more ribosomal mRNA to maintain a positive free ribosome concentration. This is consistent with the observation in (27) according to which the ratio of ribosomal promoter activity (relative to other promoters) increases with growth rate. This result implies that the correlation between ribosome production relative to other cellular proteins and the growth rate is not simply that cells with more ribosomes can grow faster, but in fact, as the growth rate increases, more *mRT* is required to maintain a stable positive steady state.

In general, the RBS strength for different genes will be different. We have also analyzed a model using different effective dissociation constant *KD1* for ribosomal and cellular mRNA and *K*_*D2*_ for the synthetic mRNA. In this case, the equations for the steepness of ribosome decline or the bifurcation point are not as simple, however numerical solutions show that the bifurcation point depends on the ratio of ribosomal mRNA to total mRNA (see Fig S7), as is the case for *K*_*D1*_=*K*_*D2*_=*K*_*D*_ as shown in Eq (2.4). Additionally, the difference between the two *K*_*D*_s determine the steepness of ribosome decline as shown in Fig S4. It is also interesting to note that for *K*_*D1*_=*K*_*D2*_, a transcritical bifurcation occurs, while when the two dissociation constants are distinct (*K*_*D1*_ ≠ *K*_*D2*_), the system passes through a transcritical bifurcation followed by a pitchfork bifurcation (Fig 4). In each case, there are additional steady state solutions for *R*<*0*, but the negative steady states are biologically irrelevant. Fig S6 shows that the main results still hold in the models where*K*_*D1*_ ≠ *K*_*D2*_: the model with constant ribosome production has no bifurcation and *R* drops slowly as *mPT* is increased; in the autocatalytic model, *R* drops much faster and *R* eventually passes through a bifurcation. Fig S6 also shows that the bifurcation in this model with *K*_*D1*_ ≠ *K*_*D2*_ also still occurs within the expression level of a high copy number plasmid. Fig S6 shows that the bifurcation point depends on the ratio between *mRT* and total mRNA, just like in the *K*_*D1*_=*K*_*D2*_=*K*_*D*_ case.

**Figure 4.**
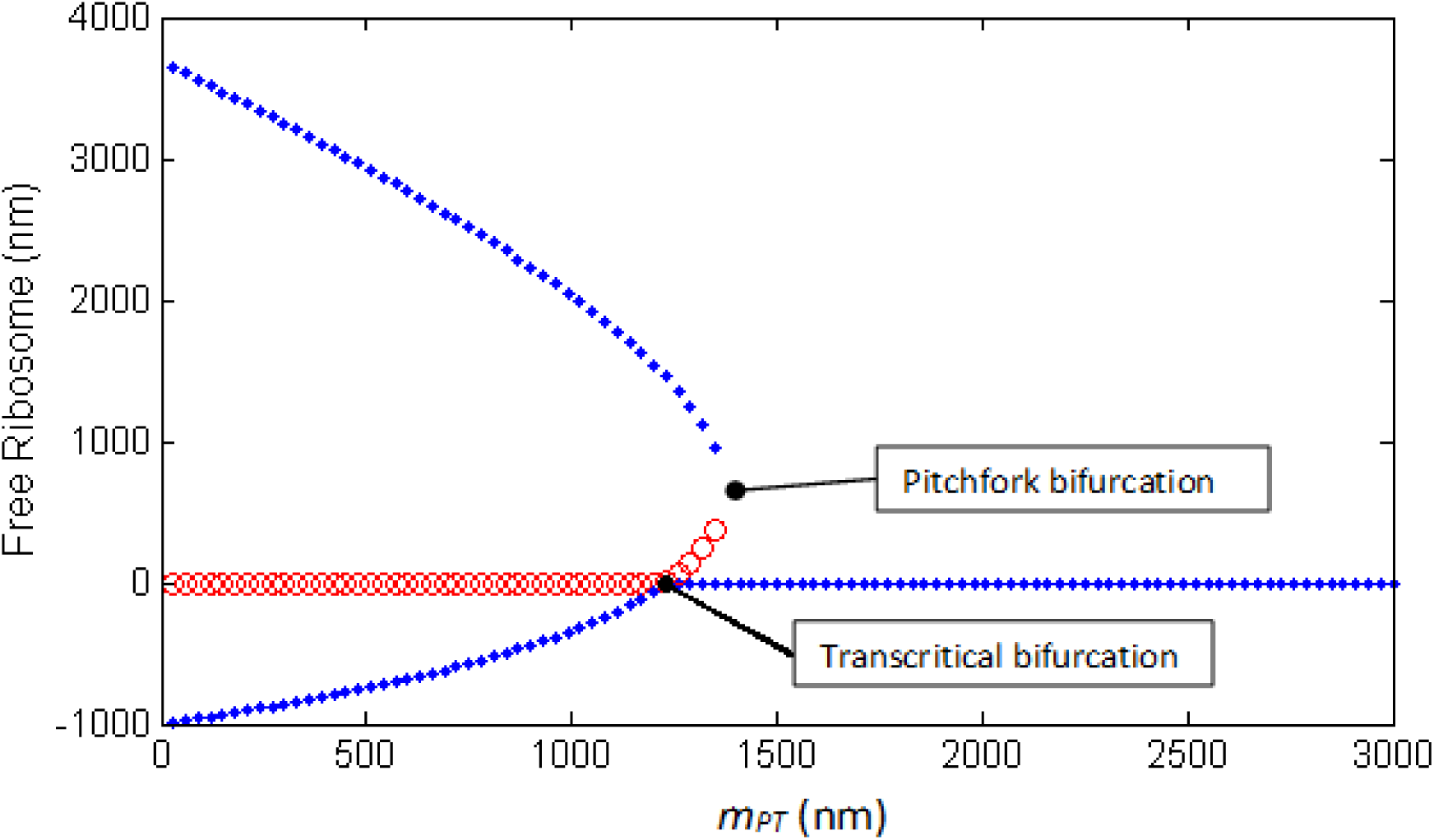
Bifurcation diagram for autocatalytic model when *K*_*D1*_≠*K*_*D2*_. Here, the synthetic gene has stronger RBS (*K*_*D1*_=2000 nm, *K*_*D2*_= 1000 nm). Parameter values other than *mPT* are as given in Fig 2. At low *mPT*, there are two stable steady states, one at *R*>*0* and one at *R*<*0* (which is physiologically irrelevant), and an unstable steady state at *R*=*0*. As *mPT* is increased, this system passes through a transcritical bifurcation in which the *R*=*0* equilibrium point becomes stable. At this point we have two equilibrium points at *R*>*0*, one of which is stable. As *mPT* is further increased, there is a pitchfork bifurcation and the three steady states collapse into one stable steady state at *R*=*0*.

### C. Model with ribosome autocatalysis and negative feedback

Regulation via ppGpp depends on the concentrations of free ribosome and the proteins RelA and DksA. Since RelA and DksA are translated by ribosomes, this feedback ultimately depends on the free ribosome concentration. In the third model, thus we incorporate negative feedback regulation on ribosome production, where ribosome production is inhibited by free ribosome concentration, modeled below using the Michaelis-Menten enzyme inhibition form:

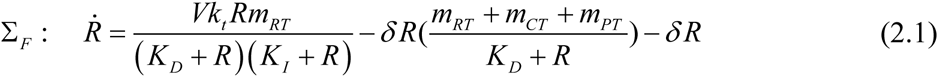

where *K*_*I*_ is the inhibition constant and *V* is a normalizing constant to maintain the same *R(mPT*=*0)* steady state as Σ_A_. Other parameter values and the references are given in Table 1.

Solving for the steady state concentration of *R* we find three possible solutions. The third solution is negative, while the non-negative solutions are given by (see SI Section E):

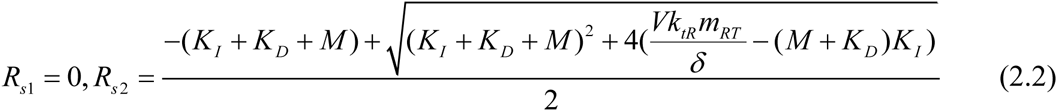

For low *mPT*, the *R*=*0* equilibrium point is unstable while the other two equilibria are stable, however physiologically the *R*<*0* steady state is irrelevant (see Fig S3). As *mPT* is increased, the system passes through a transcritical bifurcation where the *R*=*0* equilibrium point becomes stable and we have two *R*<*0* equilibria, one of which is stable and the other unstable.

The change in free ribosome concentration is given by (see SI Section E):

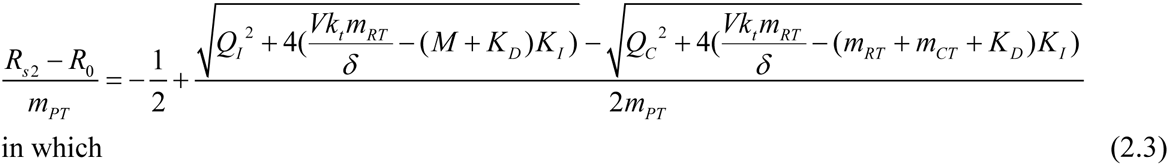

*Q*_*I*_ = *K*_*I*_ + *K*_*D*_ + *M*, *Q*_*C*_ = *K*_*I*_ + *K*_*D*_ + *m*_*RT*_ + *m*_*CT*_

When negative feedback is introduced, the drop in ribosome concentration can reach similar levels as in the non-autocatalytic model with constant ribosome production, depending on the feedback strength. When the feedback is strong enough (*K*_*I*_ is low), the autocatalytic system with feedback can maintain steady state ribosome concentrations above the constant production system for any *mPT* level, for example when *K*_*I*_=1000-2000 nm as shown in Fig 5. In this case, the second term on the right hand side of equation (3.3) is always positive, therefore the drop in ribosome concentration is again bounded below by a line with slope of −0.5. However, when 2*m*_*PT*_*K*_*I*_ > *m*_*PT*_^2^ + 2*m*_*PT*_(*K*_*D*_ + *m*_*RT*_ + *m*_*CT*_), the second term on the right hand side of Eq. (3.3) is no longer positive and ribosome concentration can drop below the lower bound of the constant production case. The free ribosome level drops faster with increasing demand when the feedback is weak (*K*_*I*_ is high), as shown in Fig 5. The feedback must therefore be strong enough to maintain robustness of the steady state ribosome concentration to changes in demand.

**Figure 5.**
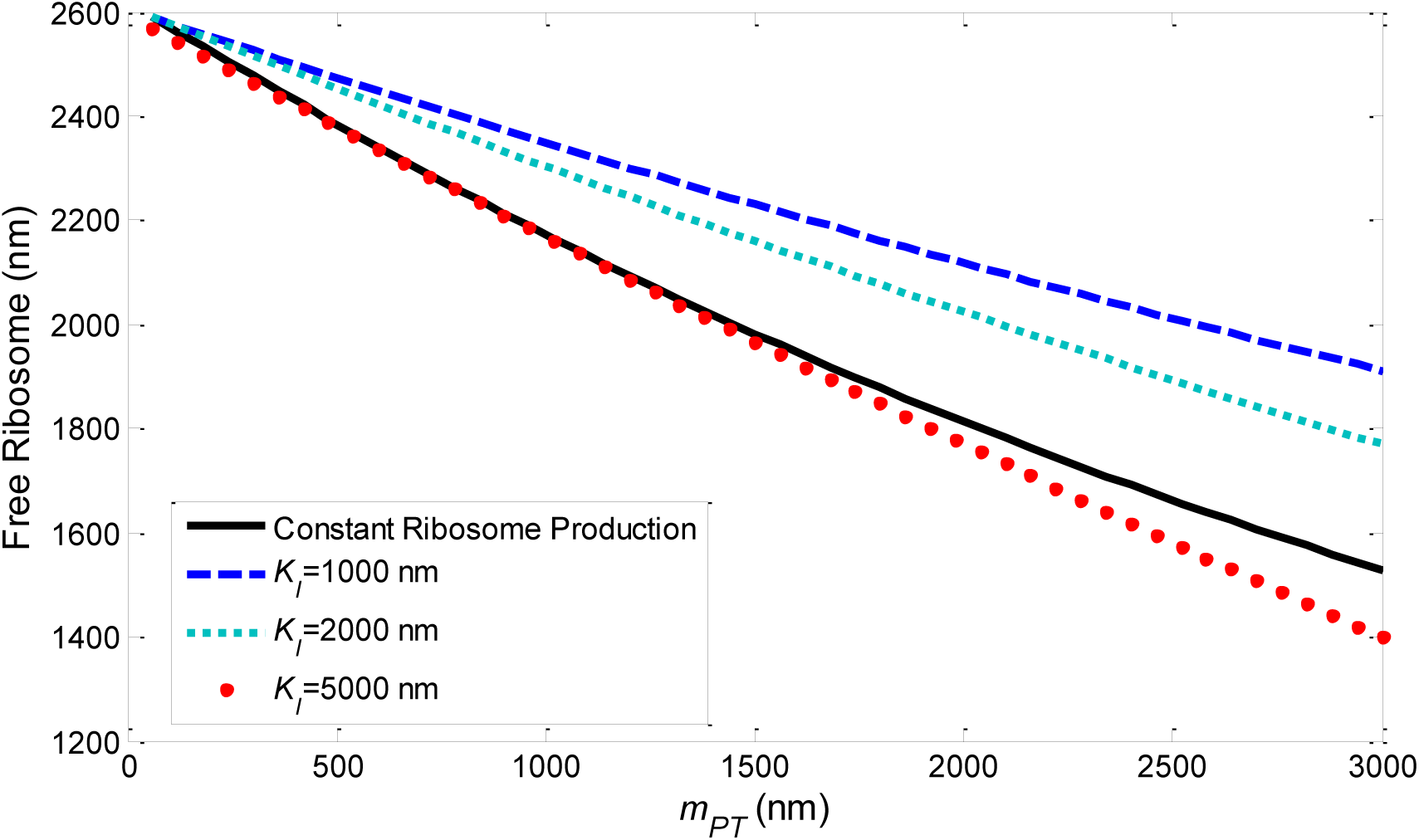
The change in free ribosome level with increasing demand for various inhibition constant *(KI)*. Higher *KI* indicates weaker feedback and results in a less robust response: ribosome level drops faster when *KI* is high. *mRT*=*550* nm, *mCT*=*5500* nm, *k*_*t*_=17.61 hr^−1^, *K*_*D*_ =1000 nm, *3*=*1* hr-^1^. For *KI*=1000 nm, *V*=3594.58 nm. For *K*_*I*_=2000 nm, *V*=4599.66. For *K*_*I*_=5000 nm, *V*=7563.884 nm.

It is important to note that the negative feedback does not remove the bifurcation, but simply shifts the bifurcation point to a higher demand level. When there is negative feedback, a positive non-zero steady state exists when (SI Section D):

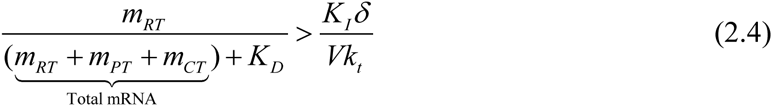

Note that since we set *V* so that the total ribosome level at zero demand is the same as in the other two models Σ_N_ and Σ_A_ (*R*_*T0*_=*7000*), *V* is large. In both cases we see from equation (3.4) that it is the ratio between ribosomal mRNA *(mRT)* over the total mRNA (plus the effective dissociation constant) that determines whether there is a positive steady state. Taking *K*_*D1*_ ≠ *K*_*D2*_ shifts the bifurcation just as we see in the system with no feedback, as shown in Fig S5, however the main results still hold (see Fig S6). When *K*_*D2*_ < *K*_*D1*_, this indicates that the RBS for the induced gene is stronger, and *R* drops faster and the system goes through bifurcation to *R*=*0* at lower demand, which is consistent with the results in (1).

### D. Temporal Dynamics

Our previous analytical and simulation results only pertain to the steady state behavior of ribosome concentration, however the dynamics of the system are equally important. In (1), the authors designed a plasmid construct consisting of an inducible fluorescent protein and a constitutive reporter protein. The authors note overshoots in the time responses of the induced gene at high induction. To simulate the temporal response of the induced gene, we add a second species to our models: a protein *P*, which is translated at maximal rate *ktP* by *R* from synthetic mRNA, *mPT. P* is also diluted with rate δ. To capture induction of *P*, we increase the value of *mPT*. Note that the differential equation for *P* is the same for all three systems Σ_N_, Σ_A_, and Σ_F_.

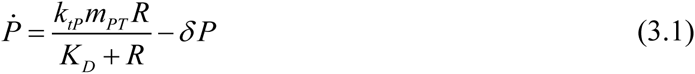

It is important to note the following observations from (1): 1) the overshoot is only seen in the induced gene (*P*), while both ribosomal and other endogenous genes display a smooth response with no transients, and 2) the overshoot does not occur at lower demand but emerges as the demand/induction level is increased.

Simulations of Σ_A_ show that there can indeed be an overshoot in the expression of the induced gene *P* when *mPT* is high enough; there is no overshoot in the free ribosome level (Fig 6B), which is consistent with the experimental observations in (1). The difference in transient responses between *P* and *R* can be explained using control theory and is due to the distinct transfer functions obtained by taking *P* or *R* as the system output, as detailed in SI Section H. As *mPT* is increased even higher, there is a spike in *P* before both *P* and *R* drops to zero. This overshoot only occurs in the autocatalytic system, while the system with no autocatalysis displays no overshoot (Fig 6A), indicating that incorporating autocatalysis in the model is important to capture the dynamics of the system’s response.

**Figure 6.**
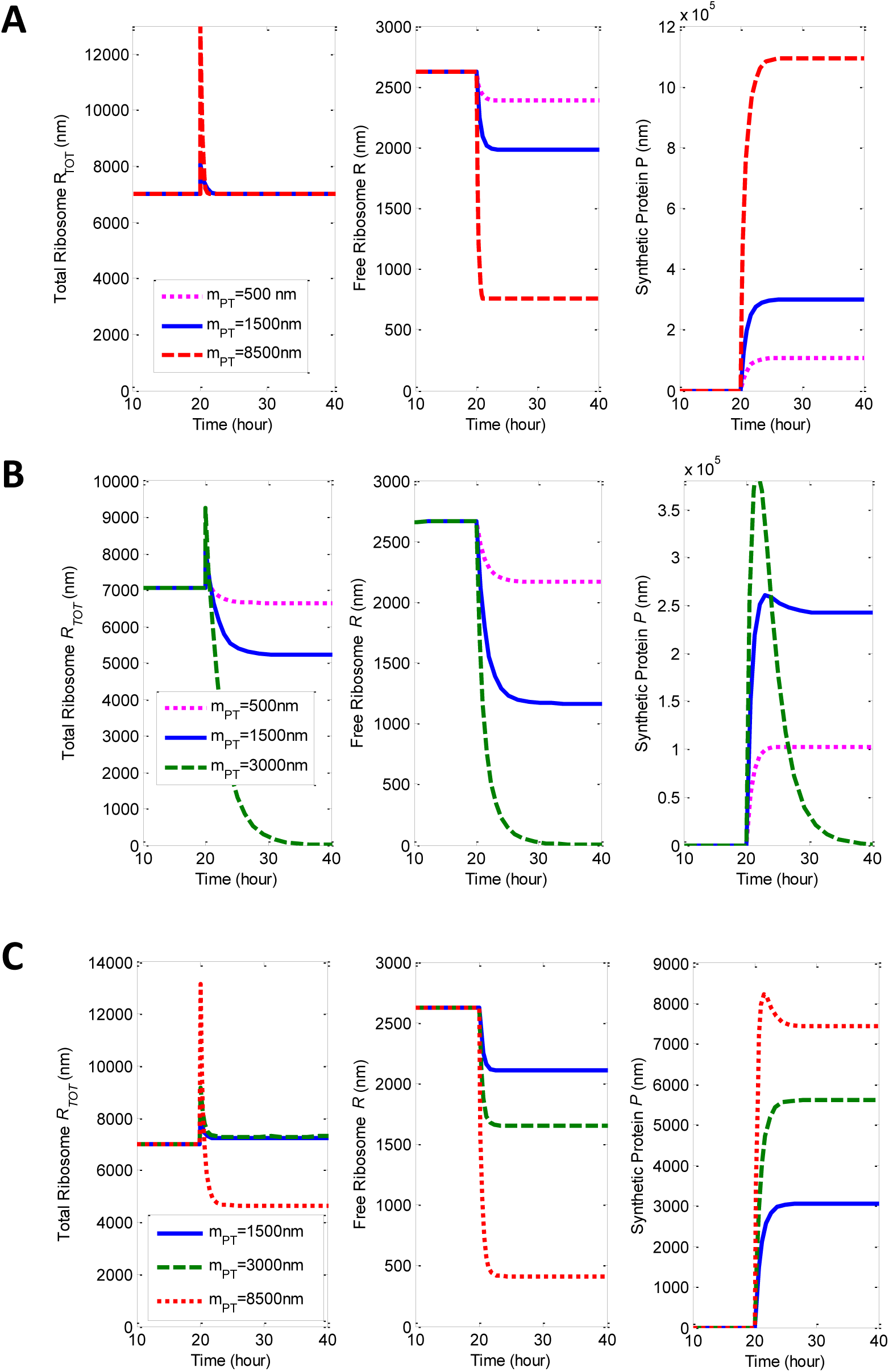
Temporal responses of the total ribosome level (*R*_*TOT*_, left), free ribosome level (*R*, middle), and expression of induced gene (*P* from Eq (2.5), right) when *mPT* is applied at time=20 hours. A) Temporal responses of the system with constant ribosome production. B) Temporal responses of the autocatalytic system without feedback. C) Temporal responses of the autocatalytic system with negative feedback. As we increase the amount of demand applied, we see an overshoot in the expression of the synthetic protein and the total ribosome level, but no transient in the free ribosome levels. The overshoot in the feedback system occurs at a higher demand than without feedback, where the response drops to a steady state of *R*=*0* and *P=0. K*_*I*_=3000 nm, *V*_*max*_=7000 nm, *m*_*RT*_=550 nm, *m*_*CT*_=5500 nm, *k*_*t*_=17.61 hr^−1^, *k*_*tP*_=300 hr^−1^, *K*_*D*_=1000 nm, δ=1 hr^−1^. For the system with feedback, *V*=5621.8 nm.

At the experimentally relevant range of demand (achievable by expressing synthetic genes on a high copy number plasmid), the steady state response of the autocatalytic system with strong feedback can be quite similar to the case with constant ribosome, which is consistent with the fact that previously published models using constant ribosome pool predicts the resource competition behavior quite well. However, the two systems can display very different behavior in the transient response. The autocatalytic system with feedback still displays overshoot in *P* when the demand is high enough as shown in the time response plots in Fig 6C. This overshoot occurs only in the autocatalytic system with and without feedback, but no overshoot occurs in the non-autocatalytic system, no matter how high the demand is (Fig 6A). The simulations shown in Fig 6 are from the nonlinear model, but we can explain the emergence of the overshoots by analyzing the linearized systems using control theoretic tools (see SI Section H). Additionally, to validate the results from the reduced model, we simulate the time responses for the non-reduced models, which display the same results (see Figs S12, S13, and S14 in SI Section I).

## Discussion

In this paper, we explored the effects of autocatalysis and feedback in ribosome production in terms of resource competition and system robustness in both steady state and transient responses. We showed that autocatalysis in ribosome production results in a bifurcation where beyond a certain level of demand, the only positive steady state is at *R*=*0*. In this condition, the cell is depleted of ribosomes, thus no protein translation can occur, resulting in cell death. Autocatalysis thus greatly affects the robustness of the system at steady state. However, the detrimental effects of autocatalysis can be mitigated by negative feedback. The feedback regulation does not remove the bifurcation, however, but simply shifts it to a higher demand level. A strong enough negative feedback can shift the bifurcation point to a demand level that is physiologically unachievable, therefore guaranteeing robustness for physiologically relevant values of mRNA levels. The negative feedback also leads to an increase in total ribosome concentration as we increase the demand, which is consistent with the hypothesis presented in (11), where the authors measure the cost of unneeded proteins in the cell. In (11), Shachrai et al found that this cost is reduced after several generations of exponential growth and hypothesize that it is due to the correction in ribosome levels to compensate for increased translation demand. From an evolutionary point of view, autocatalysis in ribosome production may be unavoidable as protein subunits may be required for better functionality and stability of the ribosome complex. Therefore, the negative feedback may have evolved to compensate for the detrimental effects of this necessary autocatalytic mechanism.

Some simplifying assumptions were made in our models to facilitate analysis. We have assumed that production of ribosomes follow a first order Hill function, however we show in SI Section C that the results also hold for higher order Hill functions. As discussed above, we also simplified the model by assuming the same ribosome dissociation constant (*KD*) for all mRNAs, however in SI Section F we show that the general results hold for distinct RBS strengths (different *KD*s) for different mRNA types.

We have shown a relationship between the ratio of ribosomal mRNA over total mRNA to the bifurcation point, indicating that higher ribosome production leads to higher robustness. We can also explain the correlation between the ratio of ribosome production over other cellular proteins and the growth rate where higher growth rate requires higher ribosome production to maintain robustness. Even though our model does not include the effects of ribosome concentration on growth rate (and vice versa), it shows that higher ribosome concentration is necessary to achieve stability at higher growth rates.

## Conclusion

While negative feedback can recover the steady state responses of the autocatalytic system to the level seen in the system with constant ribosome production, autocatalysis still has a notable effect on the temporal dynamics. In particular, the overshoots seen in the experimental results presented in (1) can only be explained by the autocatalytic models, with or without feedback. Depending on the synthetic biology application, such an overshoot may be important for the circuit functionality and therefore autocatalysis should be included in the models.

## Methods

Simulations were performed using MATLAB and Simulink. Plots were produced using MATLAB.

## Acknowledgments

We thank Yili Qian for helpful discussion and comments that greatly improve the manuscript. This work was supported by the AFOSR grant FA9550-14-1-0060 and the NIH grant P50 GM098792.

## List of Supplementary Information

A. Model reduction

B. Total ribosome concentration

C. Model with Hill coefficient *n*=*2*

D. Analytical solution for bifurcation point for autocatalytic model with *K*_*D1*_=*K*_*D2*_

E. Calculating the steepness of ribosome decrease

F. Numerical solution for bifurcation point for model with 2 distinct *K*_*D*_s

G. Change in robustness as the effective inhibition strength is varied H. Dominant zeros and overshoots in the autocatalytic models

I. Transient Responses of the Unreduced Models

